# High accuracy measurements of nanometer-scale distances between fluorophores at the single-molecule level

**DOI:** 10.1101/234740

**Authors:** Stefan Niekamp, Jongmin Sung, Walter Huynh, Gira Bhabha, Ronald D. Vale, Nico Stuurman

**Affiliations:** Department of Cellular and Molecular Pharmacology and the Howard Hughes Medical Institute, University of California, San Francisco, 600 16th Street, San Francisco, CA 94158; Skirball Institute of Biomolecular Medicine, New York University School of Medicine, New York, NY 10016

## Abstract

To uncover the mechanisms of molecular machines it is useful to probe their structural conformations. Single-molecule Förster resonance energy transfer (smFRET) is a powerful tool for measuring intra-molecular shape changes of single-molecules, but is confined to distances of 2-8 nm. Current super-resolution measurements are error prone at <25 nm. Thus, reliable high-throughput distance information between 8-25 nm is currently difficult to achieve. Here, we describe methods that utilize information about localization and imaging errors to measure distances between two different color fluorophores with ∼1 nm accuracy at any distance >2 nm, using a standard TIRF microscope and open-source software. We applied our two-color localization method to uncover a ∼4 nm conformational change in the “stalk” of the motor protein dynein, revealing unexpected flexibility in this antiparallel coiled-coil domain. These new methods enable high-accuracy distance measurements of single-molecules that can be used over a wide range of length scales.

## Introduction

Understanding the spatial arrangement of biological macromolecules is crucial for elucidating molecular mechanisms. While three-dimensional structures provide insight into the mechanism of a protein, the static state alone is often insufficient to understand how macromolecular machines perform action. By labeling single-molecules or complexes at defined sites with fluorescent dyes, it is possible to obtain static or dynamic distance measurements that provide information about conformational changes or molecular interactions.

A widely used method for obtaining such distance information is single-molecule Förster resonance energy transfer (smFRET) (1) between two different colored fluorophores. However, smFRET is limited to a narrow distance range, typically 2-8 nm. The calculation of absolute distances is influenced by the orientation and chemical environment of the fluorophores (2), which are difficult to measure, and hence smFRET is most widely used to detect relative distance changes. Direct fluorescent-based measurements of longer distances can be achieved by single-molecule localization microscopy (3–5) but distances below ∼25 nm have proven to be very difficult to measure correctly. Thus, there is an existing gap in resolution (Fig. 1a) that is important to fill since it corresponds to the size distribution of many proteins and protein complexes.

**Fig. 1.**
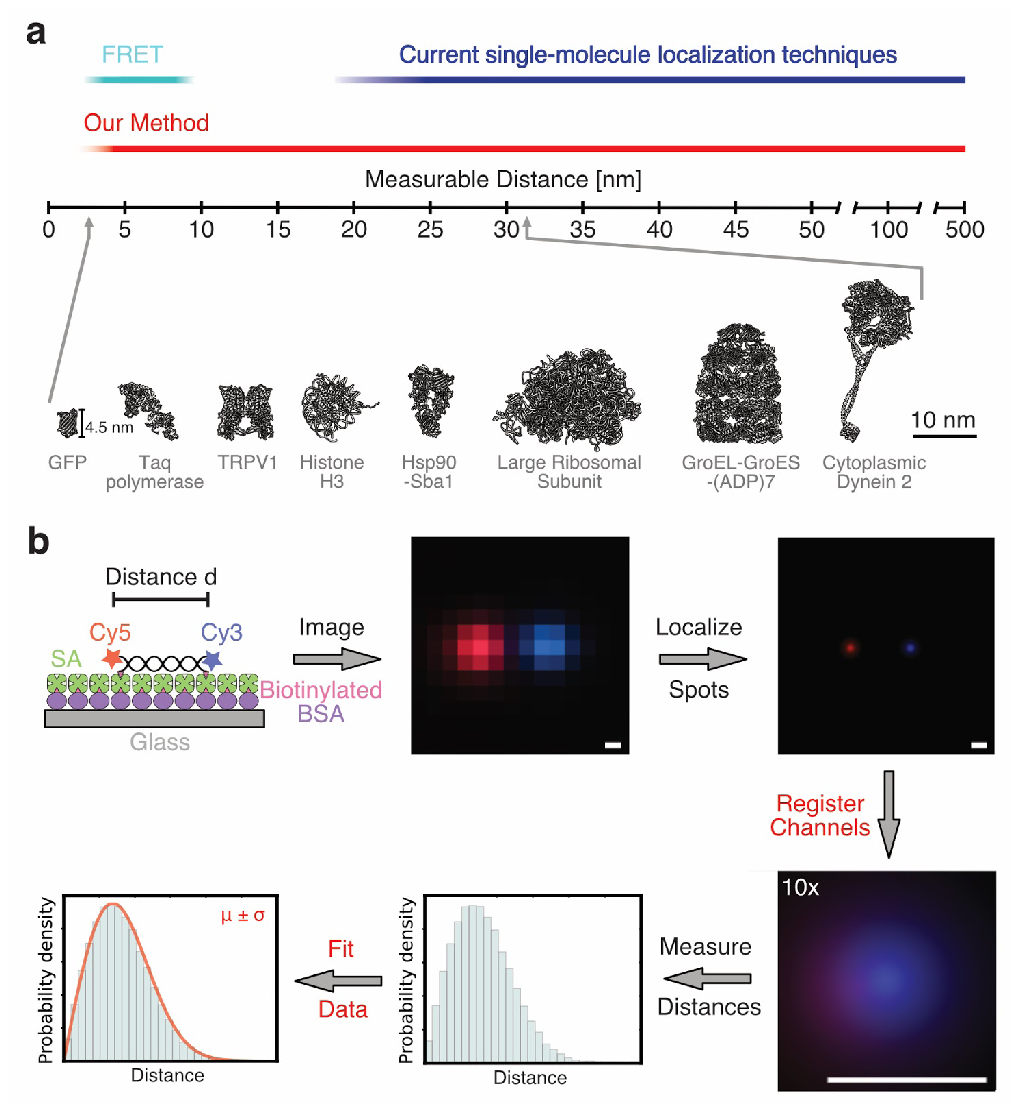
Relevance and workflow of fluorescent single-molecule distance measurements. (a) Comparison of resolution of various methods for fluorescent single-molecule distance measurements (top). Size distribution of protein structures (bottom - PDB codes from left to right: 1gfl (39), 1taq (40), 5irz (41), 1aoi (42), 2cg9 (43), 1jj2 (44), 1aon (45), 4rh7 (30)). (b) Workflow for two-color distance measurements. First, the sample of interest is labeled at specific sites with two fluorescent dyes, immobilized via biotin-streptavidin onto a glass coverslip and imaged with a TIRF microscope. Then, the exact positions of the fluorophores are determined and the positions of both dyes are registered (aligned) utilizing a registration map that was previously determined. Subsequently, the distances of all spot pairs are measured and the average distance between fluorophores is determined using a fit of a probability distribution function to the data.

Previously studies have made considerable progress in tackling distance measurements between 8-25 nm. Single-molecule high resolution colocalization (SHREC) (3, 6) resolves nanometer distances by accounting for localization errors when measuring the separation between two different color fluorophores. Pertsinidis et al. (7) developed a feedback-controlled system that enabled distance measurements with subnanometer precision, and Mortensen et al. (8) reported ∼1 nm resolution by imaging the same single-molecules multiple times. However, distance measurements with nanometer accuracy and precision have not been more broadly adopted, either because these available methods suffer from inaccuracy and/or low throughput or involve highly specialized optical setups (7).

Here, we report new methods capable of reliably measuring two-color fluorophore distances at ∼1 nm accuracy over a wide range of distances (from ∼2 nm to hundreds of nanometers) using readily available microscope hardware. To achieve this level of accuracy, we first correct for chromatic aberrations and distortions using piecewise affine transformation (9), yielding registration errors (image alignment of different fluorophores) of less than 1 nm over the entire field of view of a standard total internal reflection fluorescence (TIRF) microscope. We show that existing distance analysis methods, like those of Churchman et al. (6), are unable to measure nanometer distances correctly when the true distance and localization errors of the individual probes are similar, which is common for distances of ∼2-30 nm. To overcome these limitations, we developed two related methods: Sigma-P2D, which incorporates information about localization and imaging errors, and Vector-P2D, which makes use of averaging multiple observations of the same molecule. We applied our new methods to investigate nucleotide dependent conformational changes of the molecular motor dynein (10–12) and found that the stalk of dynein likely undergoes a large conformational change during its hydrolysis cycle (13). These results could not have been obtained by smFRET or other direct two-color imaging methods, since the distances measured changed from ∼16 nm to ∼20 nm in different nucleotide states. Thus, the two methods presented here, together with our improved image registration procedure, should have broad applications for inter- and intramolecular distance measurements, particularly in the range of 8-25 nm where techniques for two-color imaging are lacking. Our methods also are easily implemented using commercially available microscopes and open-source μManager (14) software.

## Results

### Registration error in subnanometer range

To achieve highly accurate distance measurements between two fluorophores that emit at different wavelengths, multiple obstacles have to be overcome. First, the sample of interest needs to be fluorescently labeled at specific sites and immobilized to the coverslip surface at a defined orientation (Fig. 1b). Then, one needs to image two channels, localize the individual probes, align the two channels (image registration), and calculate the distance between centroids from multiple observations of the same or multiple particles (Fig. 1b). While localization of individual fluorophores by fitting a point spread function (PSF) or 2D Gaussian to the fluorophore’s intensity distribution has been well established and delivers precision close to the theoretical limit (15), current image registration methods correct poorly for commonly observed chromatic aberrations over the entire field of view at the nanometer scale (7) or have problems in throughput since they are limited to imaging one pixel at time (8). Thus, in order to enable high-throughput and accurate two-color distance measurements, we first set out to improve two channel image registration over the entire field of view.

As multicolor fiducial markers, we imaged TetraSpeck™ beads and used a registration function to correct for the offset between color positions (Fig. 2a). While previously described registration methods either use a second-degree polynomial fit (16) or linear mapping functions (7) to calculate a registration map, we used two-step affine based registration procedure (9) commonly employed in other fields (9), but to our knowledge, not previously used to align multi-color single-molecule images. To this end, we first performed a global affine transformation to bring single spots (imaged on two different cameras) in proximity for automated pair assignment (Fig. 2b). Next, we applied a piecewise affine transformation, correcting spot positions locally (as detailed below) only using nearby fiducial points (Fig. 2b, Fig. S1). In practice, we always acquired three datasets - the first was TetraSpeck beads, the second was the sample of interest, and the third was another acquisition of the TetraSpeck beads (Fig. S1). With the corrected second fiducial marker dataset, we then calculated the target registration error (TRE) (Fig. S2), determining the deviation between the markers’ x and y positions in the two channels after alignment (Fig. 2c-h). Their mean μ_x_ and μ_y_ are the registration error along the x-axis and y-axis, respectively. The registration error σ_reg_ is given by

**Fig. 2.**
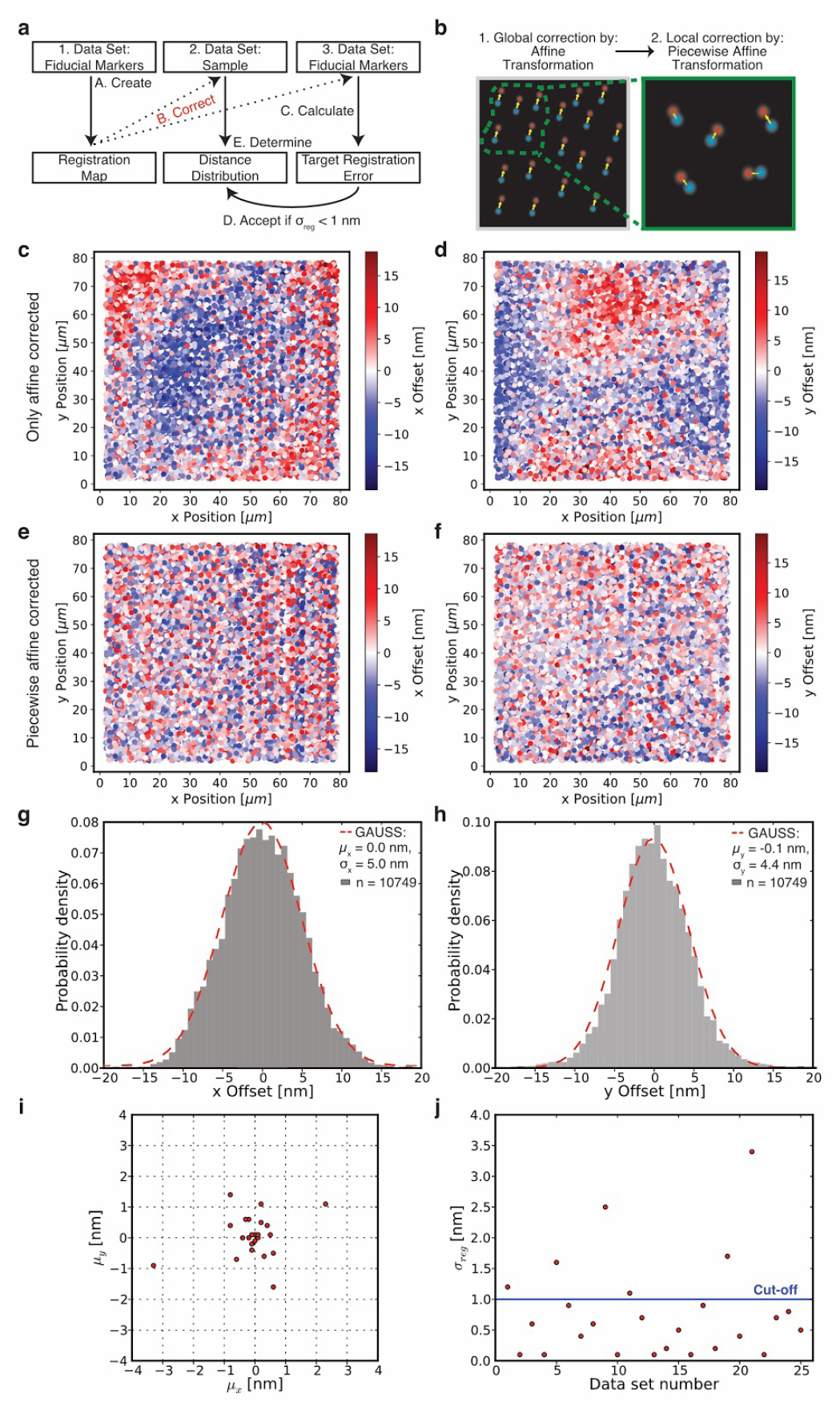
Image registration Workflow, accuracy, and reproducibility. (a) Workflow of image acquisition and registration process. (b) Procedure for image registration with affne (global) and piecewise aff.ne (local) correction. (c) Target registration error after affne correction along the x-axis. Each dot shows a single fiducial marker for which the distance offset between the two colors of the same fiducial marker is colorcoded. Negative values (blue dots) mean that channel 1 has a smaller number for its x position whereas positive values (red dots) represent fiducials where channel 2 has a smaller number for its x position. (d) Same data set as in c but the offset is along the y-axis. (e) Target registration error after piecewise aff.;ne correction along the x-axis for the same beads as in c. (f) Same data set as in e but the offset is along the y-axis. (g) Histogram of x-axis offset with Gaussian fit (dashed red line) of data shown in e. (h) Histogram of y-axis offset with Gaussian fit (dashed red line) of data shown in f. (i) X-axis μ_x_ and y-axis μy component of registration error for 25 independent image registrations. (j) Same data as in i, but registration accuracy σ_reg_ (TRE) is shown for each of the 25 data sets. We accepted data sets for distance determination if σ_reg_ < 1nm (blue line cutoff). One frame per TetraSpeck™ bead was acquired. Details of fitting parameters are provided in Table S4.

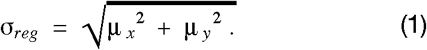

Only those samples for which σ_reg_ was < 1 nm were analyzed.

To find the optimal parameter space for image registration, we varied settings for the local piecewise affine transformation as described in more detail in the Materials and Methods section. A minimum of 10 and maximum of 100 fiducial points and a maximum distance of 2 μm resulted in optimal channel registration (Fig. S3, S4) when a sufficient number of fiducial markers was acquired. This is approximately 10,000 fiducial markers for an 80 μm x 80 μm image (Fig. S1). To obtain this number, ∼400 images with ∼25 beads per field of view were collected.

Using this approach (SI Software 1, SI Protocol), we routinely (76%) achieved registration accuracy σ_reg_ of < 1 nm (Fig. 2 i, j). Successful execution requires stable optical alignment of the two channels for the duration of the experiment (i.e. < 1 nm change in approximately 5-20 min), a high quality autofocus system (since registration changes with focus), a motorized xy-stage, minimal sample movement during image exposure (i.e. < 1 nm sample movement for approximately 1 sec), and imaging of fiducial markers for image registration and sample of interest on the same slide (Fig. S1). We noticed that the precision (σ_x_, σ_y_) for registering TetraSpeck™ beads (Fig. 2g, h) is lower than expected based on their localization errors. We found this to be caused by displacement of the color centers of TetraSpeck™ beads by a few nanometers, as reported by others (8) (SI Note 1, Fig. S5-S7). Taken together, piecewise affine alignment enables image registration at subnanometer accuracy over the entire field of view.

### Measuring distances of uniform samples

Next, we set out to optimize the accuracy and throughput of direct distance determination. Previously, Churchman et al. (3, 6) showed that distances on the scale of the localization error are non-Gaussian distributed (Fig. 3a, Fig. S8a) and described by the following two-dimensional probability distribution (P2D) (6)

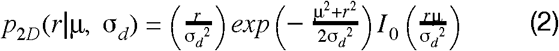

in which r is the measured Euclidean distance of individual particles, μ the estimated average distance, σ_d_ the distance uncertainty, and I_0_ the modified Bessel function of integer order zero. We refer to the true sample distance as “d”. Churchman et al. (3, 6) fit this distribution (P2D - Eq. 2) to Euclidean distance data by means of a maximum likelihood estimation (MLE) with two parameters (μ and σ_d_). We refer to this method simply as two-dimensional probability distribution “P2D”. However, using both experimental data and Monte Carlo simulations, we found that in case of small changes in distance uncertainty σ_d_, P2D yields large changes in the estimated distance μ (Fig. S8b). An approximation for σ_d_ ≥ μ shows that the probability distribution (P2D - Eq. 2) becomes independent of distance μ (SI Note 2), resulting in a fit that is driven by the distance uncertainty σ_d_. Thus, distance estimations of P2D are faulty for cases where the distance is smaller or of similar size as the distance uncertainty, which is very common for distance measurements in the range of 2-30 nm. To overcome this inaccuracy of the P2D method, we decided to fit the distance distribution with only the distance μ, and to determine the distance uncertainty σ_d_ experimentally.

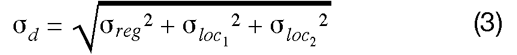

in which σ_loc_1_ and σ_loc_2_ are the localization errors of single particles of fluorophore 1 and 2, respectively, and σ_reg_ the registration error (SI Note 3). Thus, by using additional information from the images, we can fit the data only with the important parameter, the distance μ and avoid overfitting. We named this new method “Sigma-P2D” (SI Software 1, SI Protocol, Materials and Methods). Applying Sigma-P2D to Monte Carlo simulated data, for which P2D predicted an incorrect distance, we now recovered the true distance with subnanometer accuracy (Fig. 3b). Given that our new method can refine measurements made over all distances for which a distance uncertainty can be determined, we compared Sigma-P2D and P2D first using Monte Carlo simulations (SI Software 2). We generated model datasets for different ratios of distance uncertainty to distance (σ_d_ / d). We found that Sigma-P2D outperforms P2D, especially if σ_d_ ≥ μ - and that even if only 100 particles were used, Sigma-P2D estimates the true distance almost always with an error of less than 20% (Fig. 3c). Accuracy and reproducibility of Sigma-P2D can further be improved by quantifying more particles (Fig. 3c) to accuracies of better than 1% of the true distance, while P2D underestimates the distance for most conditions by almost 100%. Thus, by incorporating available knowledge of localization and registration errors we greatly improved the fitting routine and can determine distances with subnanometer accuracy and precision.

**Fig. 3.**
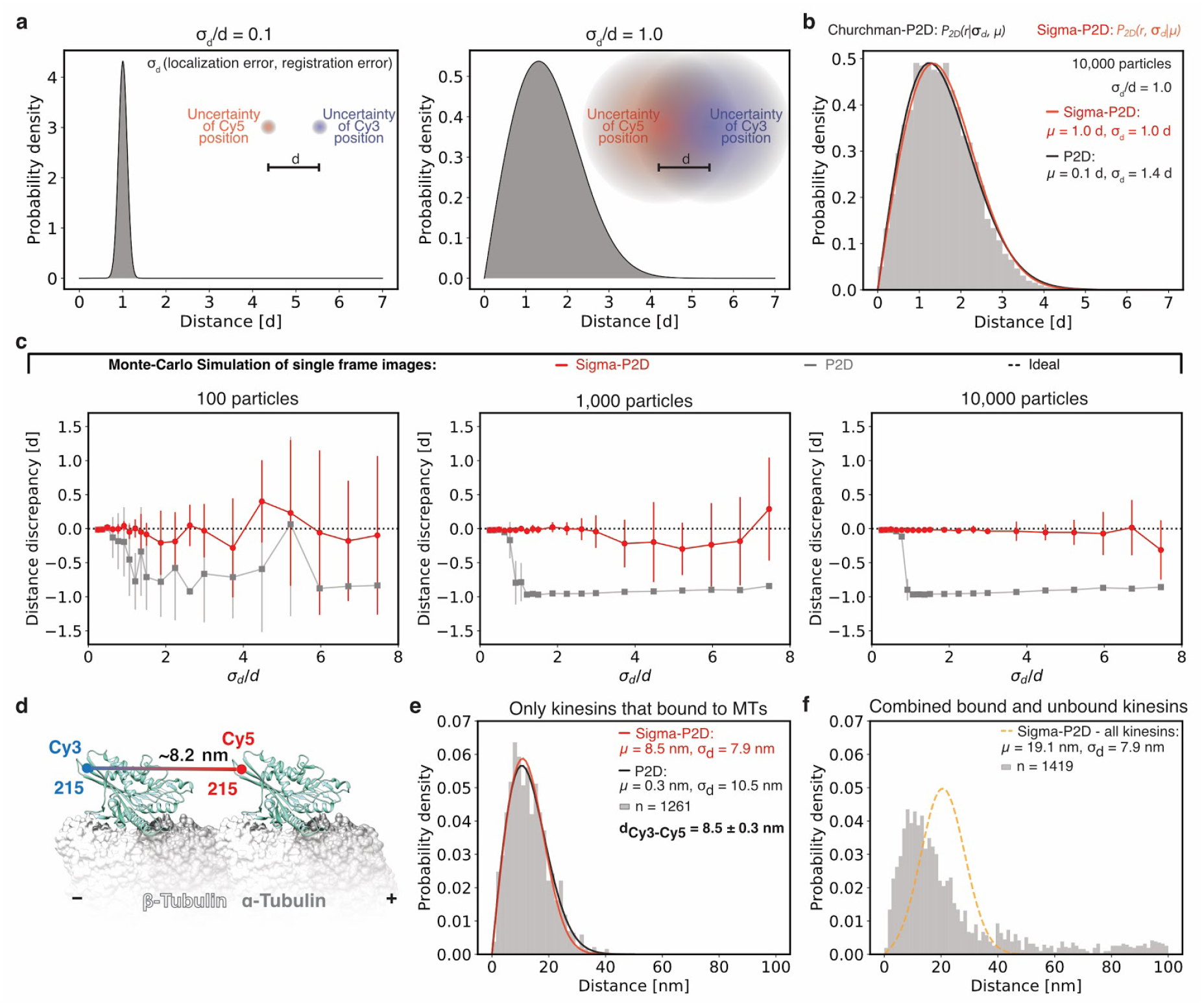
Sigma-P2D - measuring distances of uniform samples with nanometer accuracy. (a) Probability distributions of measured distances between two differently colored fluorophores separated by a true distance d for different ratios of uncertainty σ_d_ over distance d. For example, a distance uncertainty of 1 nm and a true distance of 10 nm would generate data as shown on the left while a distance uncertainty of 10 nm and a true distance of 10 nm would generate data as shown on the right. (b) Histogram of Monte Carlo simulated data with a true distance d of 1 and distance uncertainty σ_d_ of 1 fitted with Sigma-P2D (red) and P2D (black). (c) Performance of distance prediction by Sigma-P2D (red) and P2D (grey) evaluated with Monte Carlo simulated data. Average distance discrepancy from the true distance was calculated based on 10 simulations for different ratios of uncertainty σ_d_ over distance d for 100, 1,000, and 10,000 particles. Error bars show standard deviations of 10 independent simulations. (d) Diagram of two-head-bound kinesin on a microtubule based on crystal structure (PDB: 4LNU) (46) created with UCSF Chimera (47). Position of Cy3 and Cy5 dye are shown as blue and red dots, respectively. (e) Histogram of head-to-head distance measurements of rigor-bound kinesin fitted with Sigma-P2D (red) and P2D (black). The standard deviation of the head-to-head distance with Sigma-P2D fit (bold font - d_Cy3-Cy5_) was calculated by evaluating the Fisher Information matrix. (f) Histogram of head-to-head distance measurements of all kinesins (microtubule bound and unbound) fitted with Sigma-P2D (orange dashed line). Details about fitting parameters are listed in Table S4. This is possible, because all parameters of the distance uncertainty σ_d_ can be measured as it is given by

To evaluate Sigma-P2D experimentally (SI Note 3, Fig. S9-S11), we imaged a kinesin-1 homodimer for which both heads were rigor-bound with the non-hydrolyzable nucleotide analogue AMPPNP to adjacent tubulin dimers along a microtubule protofilament (17) (Fig. 3d). Based on electron microscopy data (18), the distance between the two motor domains is 8.2 nm (the tubulin dimer spacing). A kinesin motor domain construct (17, 19) with a single cysteine residue (E215C), was reacted with an equimolar mixture of maleimide-Cy3 and maleimide-Cy5. Motors that contained both Cy3 and Cy5 and that bound to a biotin-streptavidin immobilized and Alexa-488 labeled microtubule were selected for two-color distance measurements.

When fitting the data for the apparent head-to-head distance of the rigor-bound kinesins with Sigma-P2D, we measured 8.5 ± 0.3 nm (Fig. 3e), which is very close to the expected distance of 8.2 nm. Fitting the same data with the P2D method shows that P2D dramatically underestimated the distance and finds 0.3 ± 1.0 nm (Fig. 3e). Unbound kinesins had variable distances causing the probability distribution fits to yield incorrect results (Fig. 3f) since Sigma-P2D does not consider conformational heterogeneity. Hence, Sigma-P2D can only fit samples that are uniform in distance unless prior knowledge about the conformational heterogeneity σ_con_ is available. Nevertheless, utilizing Sigma-P2D we measured the head-to-head distance of a kinesin dimer with subnanometer accuracy and precision.

### Measuring average distances of heterogeneous samples

Since distance measurements for heterogeneous samples with Sigma-P2D are inadequate and many proteins and protein complexes are heterogeneous in distance, we needed an additional method. To obtain meaningful population statistics for samples which are heterogeneous in distance, it is important to improve the precision with which the two-color distances of individual molecules can be measured. To do so, we collected multiple observations (frames) of the same molecule, by time-lapse imaging (Fig. 4a). Rather than directly averaging the distance in each frame, observations of the same fluorescent pair in multiple frames are combined by first averaging distances in x and y separately, and then using these to calculate the Euclidean distance of individual particles (vector distance average). As previously shown (8), this leads to more accurate distance predictions than direct frame-by-frame Euclidean distance averaging (Fig. 4a, Fig. S12a). If for example 10,000 particles are imaged and 5 observations per particle are recorded, either all 50,000 distance measurements (frame-by-frame Euclidean distance) or all 10,000 vector averaged distances can be combined. For the vector averaged distances, the distribution is more narrow (Fig. 4b) but still not perfectly gaussian distributed. Instead of fitting with a gaussian probability distribution as done in a previously developed method (8) (here named “Vector”), we noticed that the fit can further be improved using the two-dimensional probability distribution (P2D - Eq. 2) and two parameters (μ and σ_d_). Moreover, we noticed that maximum likelihood estimation (MLE) fitting often resulted in inaccurate distance determination for experimental data since it is fairly sensitive to outliers (background noise). Therefore, we fit the P2D function by means of non-linear least squares (NLLSQ), which is more robust to background noise than MLE (see Materials and Methods). We call this method “Vector-P2D” and found that Vector-P2D outperforms Vector for all conditions tested using Monte Carlo simulations (SI Software 2). Using Vector-P2D (SI Software 1, SI Protocol), 100 particles with 20 observations each are enough to resolve distances within 5% of the true distance (Fig. 4c). Increasing the number of particles to 1000 with 20 observations results in fitted distances that diverge less than 1% from the true distance (Fig. S12b-g). To test if Vector-P2D can determine the average distance of samples that are variable in distance, we ran Monte Carlo simulations at varying degrees of sample heterogeneity σ_con_ (standard deviation of true distances in the population). Even for cases where σ_con_ is twice as large as the true distance d, we still recovered the correct population average (Fig. S13).

**Fig. 4.**
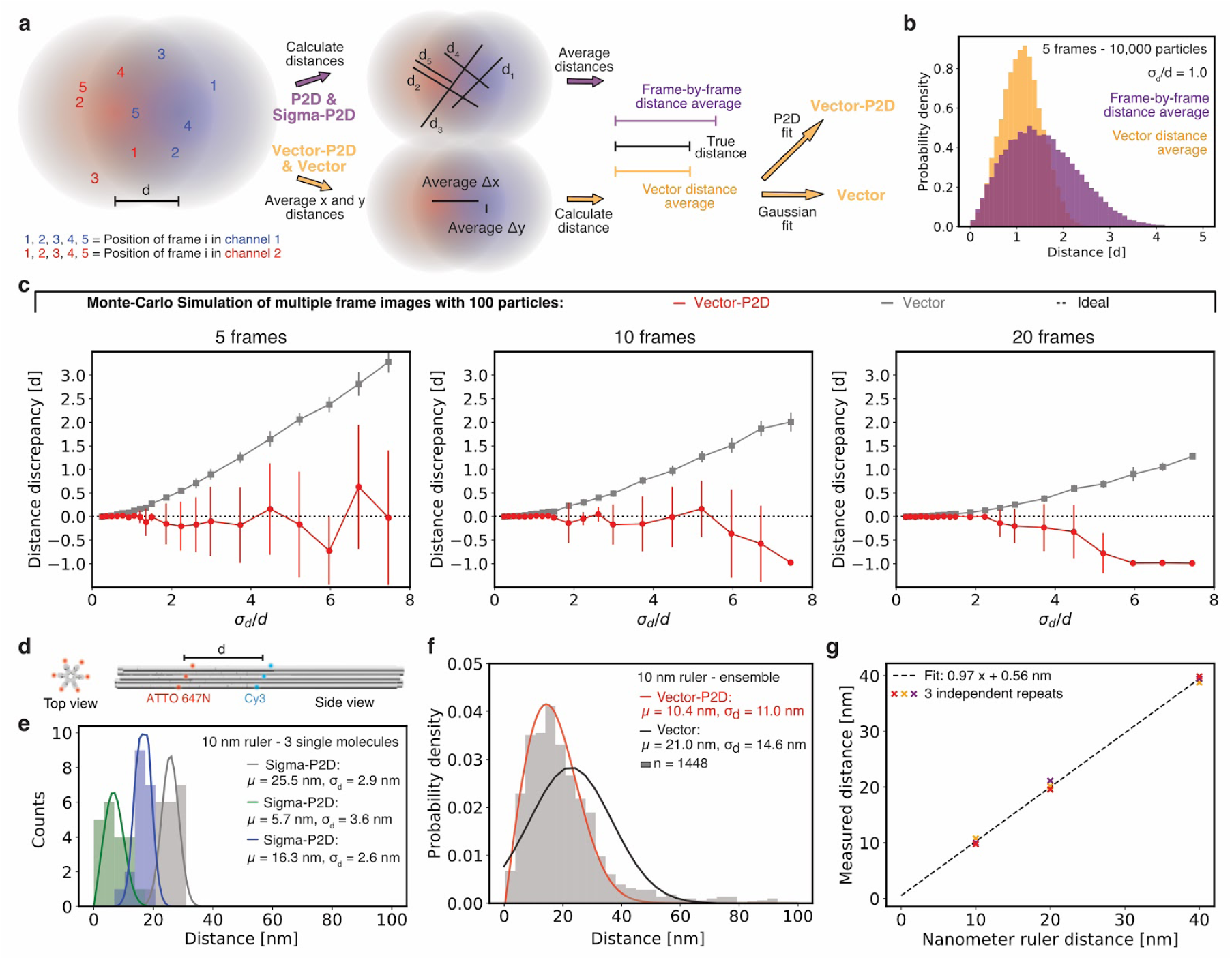
Vector-P2D - measuring distances of variable samples with nanometer accuracy. (a) Determining vector averaged distances from data with multiple observations per particle. Intensity distributions for two fluorescent molecules in a red and a blue channel at a true distance d of 1. Five independent observations of both molecules were obtained by Monte Carlo simulations (red and blue colored numbers 1 to 5). Now either the individual distances of spotpairs can be calculated first and then averaged (frame-by-frame distance average) or average distances along the x-axis and y-axis can be determined first and then used to calculate the absolute distance (vector averaged distances). The vector averaged distance distribution can then be fit with a Gaussian distribution or the two dimensional probability distribution “P2D” as shown in equation 2, which used the calculated distance μ and the distance uncertainty σ_d_ as parameters, to yield Vector or Vector-P2D, respectively. (b) Histograms for distances generated by means of Monte Carlo simulation with 5 frames (observations) per particle. Purple histogram shows the distance distribution for a frame-by-frame distance average and orange histogram shows distribution for vector averaged distances. (c) Performance of distance prediction by Vector-P2D (red) and Vector (grey) evaluated with Monte Carlo simulated data. Average discrepancy from the true distance was calculated based on 10 simulations for different ratios of uncertainty σ_d_ over distance d for 5, 10, and 20 frames. Error bars show standard deviations of 10 independent simulations. Additional data in Fig. S12. (d) Design of DNA-origami based nanorulers for which the ‘center-of-mass’ between 6-10 dyes for each of the two colors determines the distance. (e) Histogram of distance distribution of three different single-molecules of the 10 nm DNA-origami nanoruler (green, blue, and gray). Solid line is a Sigma-P2D fit. (f) Histogram of vector averaged distance measurements of multiple 10 nm DNA-origami nanorulers analyzed with Vector-P2D (red) and Vector (black). (g) Correlation between measured and expected average distance for 10, 20, and 40 nm ruler from three technical repeats. Example fits for 20 and 40 nm rulers are shown in Fig. S14. Fitting parameter details are given in Table S4.

To test the performance of Vector-P2D experimentally, we used DNA-origami based nanorulers (20, 21). The average ‘center-of-mass’ distance between Cy3 and Alexa647 fluorophore binding sites on these nanorulers is either 10 nm, 20 nm, or 40 nm. Each color has up to 10 binding sites with an expected labeling efficiency of 50-80% (Fig. 4d). Together with bleaching effects this results in variable distances of the color centers of individual rulers (Fig. 4e, Fig. S14e, f). However, when we analyzed these rulers using Vector-P2D, we found average distances that were within a nanometer of the expected values (Fig. 4f) whereas the Vector method predicted distances up to 100% larger (Fig. 4f, Fig. S14a-d). Plotting the Vector-P2D measured population distances for all three nanorulers of three repeats over the expected distances and calculating the slope, we found a slope of 0.97, very close to the ideal value of 1.0 (Fig. 4. g, Fig. S14a-d). Summarizing, using multiple observations of the same molecule and by performing a vector distance average, we can recover distances of variable samples with nanometer precision and accuracy.

### Measurements of the dynein stalk length in multiple nucleotide states

We next applied our two-color localization methods to measure conformational changes in the minus-end directed, microtubule-based motor dynein (10–12). An intriguing problem for the function of this molecular motor is the two-way communication between the catalytically active AAA ring and the microtubule binding domain through conformational changes in an intervening ∼13 nm antiparallel coiled-coil stalk (13, 22–24) (Fig. 5a). Earlier studies have suggested that local melting of the coiled-coil stalk in different states of the nucleotide hydrolysis cycle plays a major role in this communication (25–27), while others have shown that a 4 amino acid sliding between different registries is critical (13, 25). However, no direct measurements of the distances between the AAA ring and microtubule binding domain have been reported, which could help to distinguish between these models.

**Fig. 5.**
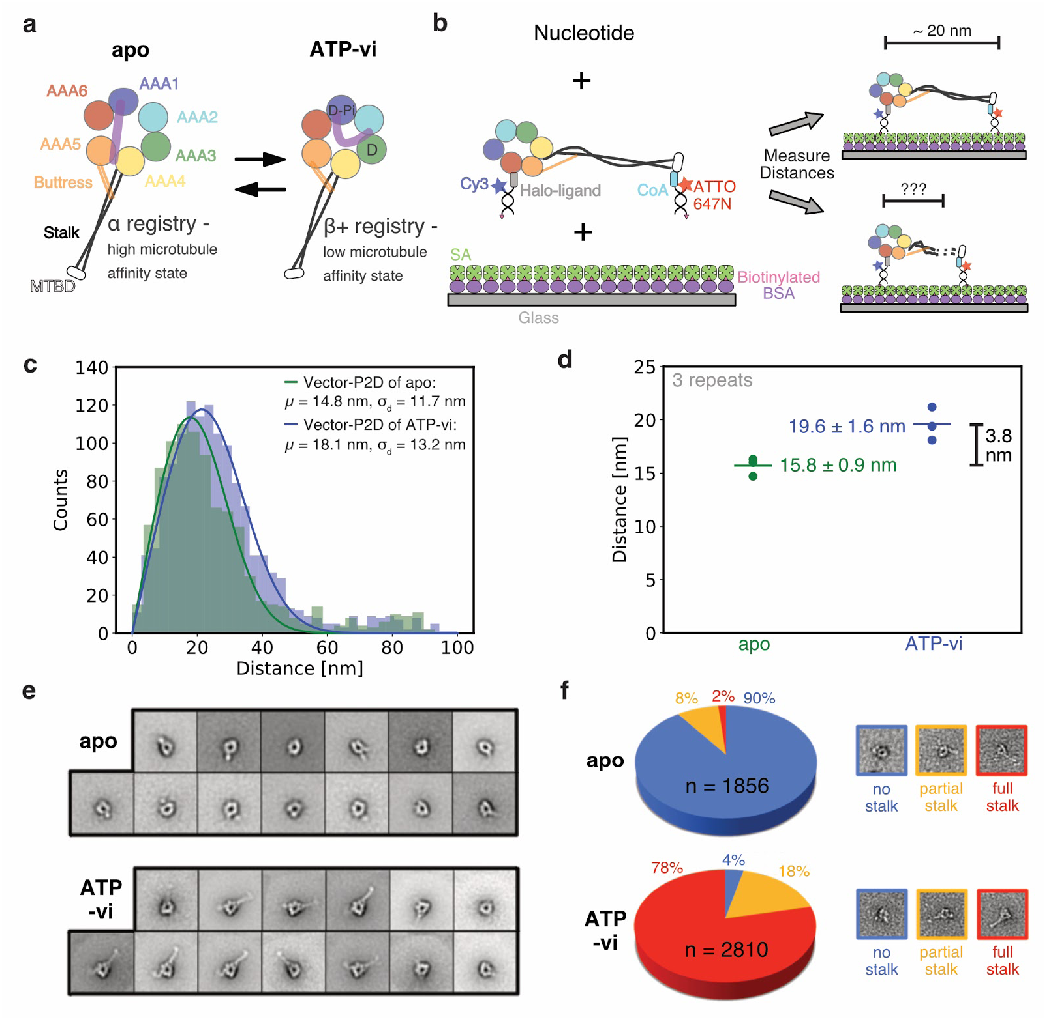
Dynein stalk conformation in two different nucleotide-bound states measured by Vector-P2D and negative stain electron microscopy. (a) Schematic of the monomeric dynein motor domain without nucleotide (apo) and bound to ATP-vanadate (ATP-vi) r ting in a high and low microtubule affinity state, respectively. Transition between both esul microtubule affinity states happens twice during the hydrolysis cycle: first detachment from microtubule by ATP binding and transition to a low affinity state and then rebinding to microtubule after ATP hydrolysis and change to a high affinity state. (b) Design for two-color fluorescent distance measurement between AAA ring and microtubule binding domain of a dynein monomer. Fluorescent dye, Halo-tag (28) or YBBR-tag (29) ligands and biotin for surface immobilization are attached to a double stranded DNA oligomer of 16 bp where Cy3 labels the Halo-tag on the C-terminus of the AAA ring and ATTO647N is attached to the YBBR tagged microtubule binding domain. If the stalk is fully extended we expect a distance of about 20 nm (30) between the two colors. (c) Histogram of vector averaged distance measurements of dynein monomer as shown in b with apo in green and ATP-vi in blue fitted with Vector-P2D. Only molecules that had both a Cy3 and ATTO647N label were selected for analysis to ensure measurement of the distance between ring and microtubule binding domain. (d) Results of distance measurements of three technical repeats of dynein monomer as shown in b with apo in green and ATP-vi in blue. Fitting was done as shown in c. Error is standard deviations of 3 technical repeats. (e) Negative stain electron microscopy class averages of dynein monomer in the apo (top) and ATP-vi (bottom) state. (f) Count and classification of individual particles from negative stain electron microscopy micrographs (as shown in Fig. S15, Table S1) into three categories (no, partial, and full stalk) for the apo state and the ATP-vi state. Single-molecule distances in c and d were obtained by selecting time-lapse series of individual molecules (see Table S6). Details about fitting parameters are provided in Table S4.

To tackle this problem, we prepared a yeast cytoplasmic dynein monomer with a C-terminal Halo-tag (28) and a YBBR-tag (29) that was inserted into the microtubule binding domain (Fig. 5b). Based on structural data, we predicted that the distance between Halo- and YBBR-tag for a fully extended stalk would be ∼20 nm (30) (Fig. 5b). To simultaneously immobilize and fluorescently label dynein, both tags were labeled with a 16 bp long double stranded DNA that was biotinylated at one end and dye-labeled at the other. We then imaged dynein in the apo and ATP-vanadate (ATP-vi) state and measured the distance between the fluorescent labels using Vector-P2D since we expected a heterogeneous distance distribution. Using three technical repeats, we measured a distance of 19.6 ± 1.6 nm for the ATP-vanadate state (Fig. 5c, d). This is consistent with the X-ray crystallographic studies (30). However, in the apo state (no ATP), we measured a distance of 15.8 ± 0.9 nm between the Halo-tag on the ring and the YBBR-tag in the microtubule binding domain. This shorter distance cannot be explained by the “simple helical sliding” model (13, 25), which predicts essentially no distance change.

To further understand the structural basis of our two-color fluorescence measurement, we turned to negative stain electron microscopy. Two-dimensional class averages for the ATP-vanadate bound state show a clear density for the stalk and microtubule binding domain in most classes (“full stalk”). In contrast, the stalk density in the apo state was rarely observed (“no stalk”) (Fig. 5e). This suggests two possibilities: 1) The angle of the stalk differs significantly in the individual molecules in the apo state, leading to these angles being averaged out in 2D classes, or, 2) The coiled-coil stalk of individual particles in the apo state cannot be identified in the micrographs, suggesting a large-scale conformational change in the stalk. To address these two possibilities, we analyzed the negative stain data on a single particle level. Individual particles for multiple nucleotide states were manually scored as belonging to one of three categories: no stalk, partial stalk, and full stalk. Consistent with the results of the class averages, we saw full stalk density for 79% of all particles in the presence of ATP-vanadate and only for 4% in the apo state (Fig 5f, Fig. S15, Table S1). Moreover, almost all particles (90%) in the apo state do not have any visible density of the stalk, whereas the number of particles for the ATP-vanadate state is a little more distributed among all three categories. This agrees well with our two-color fluorescent distance measurements as the distance distribution in the apo state is more narrow than in the ATP-vanadate state. These results suggest a disorder-to-order transition in the stalk that is important for dynein’s mechanochemical cycle. The negative stain electron microscopy data also suggest local melting or conformational changes of the stalk in the apo state, which is consistent with our two-color fluorescent distance measurements.

## Discussion

Here, we described single-molecule two-color fluorescent microscopy methods that provide nanometer accuracy distance measurements on the length scale of most macromolecules (2-30 nm). Using Monte Carlo simulations and experiments, we show that our techniques enable distance measurements from ∼2 nm to hundreds of nm and can operate with heterogeneous samples. Thus, our methods fill a resolution gap from 8 nm (upper distance of smFRET) to 25 nm (lower bound of current single-molecule localization methods). Applying our methods to the molecular motor dynein, we found that the dynein stalk likely undergoes large conformational changes in different nucleotide states.

### Distance calculations with nanometer accuracy

While smFRET can accurately determine distances in a high-throughput fashion, it is limited to distances that are <8 nm (1, 2). Furthermore, absolute distance measurements are difficult because smFRET is sensitive to fluorophore orientation, which is often assumed to be randomly oriented but non-trivial to measure. There are some existing single-molecule localization methods that can be used at the 8-25 nm range but all of these methods face certain limitations. For instance, single-molecule high resolution colocalization (SHREC) (3, 6) inaccurately determines distances for cases where distance uncertainty and distance are of similar size. We overcame this limitation by using additional experimental information from the images (Sigma-P2D). A method developed by Pertsinidis et al. (7) also achieves nanometer resolution but is limited to single pixel measurements and requires highly specialized optical setups, whereas our new methods work on the entire field of view of a standard TIRF microscope. Lastly, a method by Mortensen et al.(8) resolves nanometer distance with lower resolution (Vector method) and only measures tens of molecules whereas our methods can measure up to 10,000 molecules in a single experiment.

In general, we significantly improved and extended existing methods by using additional experimental information (Sigma-P2D) and by improving analysis techniques of multiple observations of the same particle (8) (Vector-P2D). Our Sigma-P2D approach recovers the distance from a collection of uniform particles, while Vector-P2D measures average distances of samples that are variable in distance (Fig. S16). Even though Vector-P2D also works well for samples that have a uniform distances, the Sigma-P2D method is useful to determine whether or not a sample is uniform in distance (Fig. S16). Moreover, when only one or a few frames (typically 1-5 frames) can be imaged, Sigma-P2D results in higher accuracy distance measurements than Vector-P2D (Fig. S17). This is particularly important for future extensions of our methods to dynamic measurements. Our immediate interest lies in analyzing stepping motor proteins for which only a few images can be collected in between steps. We are planning to test such uses of Sigma-P2D in the near future.

Like other existing localization-based two-color distance measurement methods (7, 8, 13, 25), our methods require surface immobilization of the sample and are limited to projections in two-dimensions. Nevertheless, using versatile labeling techniques (such as the DNA-based surface coupling combined with labeling as we used for the dynein experiment), we believe that there are many ways to obtain useful information - difficult or impossible to acquire otherwise - while being aware of this limitation. In summary, our new methods, Sigma-P2D and Vector-P2D, together with the piecewise image registration and the μManager plugin (14) allow distance measurements in less than two hours on a standard TIRF microscope, enabling high-throughput distance measurements with nanometer accuracy.

### Stalk of dynein likely undergoes large conformational changes

In order for dynein to step along microtubules, the hydrolysis state of the nucleotide binding AAA ring is coupled to microtubule affinity of the microtubule binding domain through the stalk (13, 22–24). Several studies suggest that local melting of the coiled-coil in different states of the nucleotide hydrolysis cycle plays a major role in this communication (25–27), while others have shown that sliding between different registries is essential (13, 25). However, no direct measurements of the distances between the AAA ring and microtubule binding domain have been reported. Using the Vector-P2D method, we measured this distance directly in different nucleotide states and found evidence for a large conformational change in the dynein stalk. These measurements would not have been possible with other methods such as smFRET since we could not have placed any fluorescent labels in the working range of smFRET (2-8 nm) as the stalk of dynein is 13 nm long. Moreover, the negative stain electron microscopy approach also does not allow direct distance measurements since we only get information about visible vs. not visible density.

Our observations do not rule out registry sliding of the stalk (13, 25), however, the changes in distance cannot be explained by simple sliding and small conformational rearrangements alone and thus provide evidence for a local stalk “melting” (25–27). Based on the distance measured in the apo state, we speculate that some part of the stalk between the microtubule binding domain and the buttress / stalk interaction is involved in these conformational changes. This is in good agreement with the model in which a highly conserved tryptophan in the stalk close to the buttress melts coiled-coil 1 and triggers a conformational rearrangement of the stalk (31).

Finally, the theoretical concepts and their application to nanometer distance measurements presented in this work are not limited to two-color fluorescent single-molecule localization microscopy but apply to all distance measurements where the distance is similar to the error and thus also to other super-resolution imaging techniques (32). As these methods venture into the regime of nanometer resolution (33), we anticipate that our methodology and open-source software will be useful for a broad range of super-resolution fluorescence microscopy technologies.

## Materials and Methods

### Flow-cell preparation

We custom made three-cell flow chambers using laser-cut double-sided adhesive sheets (Soles2dance, 9474-08×12 - 3M 9474LE 300LSE), glass slides (Thermo Fisher Scientific, 12-550-123), and 170 μm thick coverslips (Zeiss, 474030-9000-000). The coverslips were cleaned in a 5% v/v solution of Hellmanex III (Sigma, Z805939-1EA) at 50°C overnight and washed extensively with Milli-Q water afterwards. Flow-cells were assembled so that each chamber holds ∼10 µl (Fig. S1).

### Fluorescent beads for image registration

We used TetraSpeck™ beads (Thermo Fisher Scientific, T7279) with a diameter of ∼100 nm for image registration. To prepare the beads for imaging we added 10 µl of 1 mg/ml Poly-D-lysine (Sigma, P6407) in Milli-Q water to the flow-cell and incubated for 3 min, washed with 20 µl of BRB80 (80 mM Pipes (pH 6.8),1 mM EGTA, and 1 mM MgCl2) and then added 10 µl of 1:1000 diluted TetraSpeck™ beads in BRB80 and incubated for 5 min. Finally, we washed the flow-cell with 40 µl of BRB80.

### Preparation of dsDNA samples

For the 30 bp single biotin dsDNA construct we used

~~~
Strand A:
/5Cy3/GGGTATGGAGATTTTTAGCGGAGTGACAGC/3Cy5Sp/
strand B:
/5BiosG/AAAAAAAAAAAAGCTGTCACTCCGCTAAAAATCTCC ATACCC
~~~

both purchased from Integrated DNA Technologies (Skokie, IL). The double stranded constructs were assembled by mixing 10 µM of strand A and B with Assembly Buffer (20 mM Tris (pH 8.0), 1 mM EDTA, and 2.5 mM MgCl2) and heating the mixture to 95°C for 5 min, followed by cooling down to 20°C at a rate of 1°C per minute. For imaging, we diluted the constructs in Assembly Buffer to 3 pM. Samples for imaging are prepared by adding 10 µl of 5 mg/ml Biotin-BSA (Thermo Fisher Scientific, 29130) in BRB80 to the flow-cell, incubation for 2 min., addition of 10 µl of 5 mg/ml Biotin-BSA in BRB80, incubation for 2 min, washing with 20 µl of BRB80, addition of 10 µl of 0.5 mg/ml Streptavidin (Thermo Fisher Scientific, S888) in PBS and a 2 min incubation. We then washed with 20 µl of PBS pH 7.4, added 10 µl of 3 pM dsDNA construct in PBS pH 7.4 and incubated for 5 min. Next, we washed with 30 µl of PBS pH 7.4 and finally added the PCA/PCD/Trolox oxygen scavenging system (34) in PBS pH 7.4.

### DNA-origami standards

Custom DNA origami nanorulers (21) were purchased from GATTAquant GmbH (Braunschweig, Germany). The nanoruler design is based on the 12HB and is externally labeled with fluorescent dye molecules (Cy3 and Alexa647). The ‘center-of-mass’ between the Cy3 binding sites and the Alexa647 binding sites is either 10 nm, 20 nm, or 40 nm. Each color has up to 10 binding sites with an expected labeling efficiency of 50-80%. In addition, each nanoruler has multiple biotins attached for immobilization on a coverslip. Samples for imaging are prepared by twice adding 10 µl of 5 mg/ml Biotin-BSA in BRB80 to the flow-cell and incubation for 2 min., washing with 20 µl of BRB80, addition of 10 µl of 0.5 mg/ml Streptavidin in PBS and a 2 min incubation. We then washed with 20 µl of PBS pH 7.4 supplemented with 10 mM MgCl_2_. In a next step 10 µl of DNA-origami ruler was added and incubated for 5 min. Next, we washed with 30 µl of PBS pH 7.4 supplemented with 10 mM MgCl_2_ and finally added the PCA/PCD/Trolox oxygen scavenging system (34) in PBS pH 7.4 supplemented with 10 mM MgCl_2_.

### Kinesin cloning, purification and labeling

The kinesin construct was cloned and purified as previously described (17, 19). Briefly, cysteine residues were introduced into a ‘cysteine-light’ human kinesin-1 dimer that is 490 amino acids long (K490). The homodimeric E215C K490 contains a carboxy-terminal His_6_ tag.

The plasmid was transfected and expressed in Agilent BL21(DE3). Cells were grown in LB at 37°C until they reached 0.6 OD_280_, expression was induced by addition of 1 mM IPTG and cells were incubated overnight at 18°C. Cells were pelleted and harvested in lysis Buffer (25 mM Pipes (pH 6.8), 2 mM MgCl_2_, 250 mM NaCl, 20 mM imidazole, 2 mM TCEP, 5% sucrose), and lysed in the EmulsiFlex homogenizer (Avestin) in the presence of protease inhibitors. After a spin in a Sorvall SS-34 rotor for 30 minat 30,000 x g, the supernatant was loaded onto a Ni-NTA resin (QIAGEN, 30210) and washed with additional lysis Buffer. Then the protein was eluted by adding 300 mM of imidazole to the lysis Buffer. The elutions were dialyzed overnight against dialysis Buffer (25 mM Pipes (pH 6.8), 2 mM MgCl2, 200 mM NaCl, 1 mM EGTA, 2 mM TCEP, 10% sucrose).

Afterwards, the E215C K490 was reacted for 4 h at 4°C with Cy3-maleimide (GE Healthcare, PA13131) and Cy5-maleimide (GE Healthcare, PA15131) at a motor/Cy3 dye/Cy5 dye ratio of 1:10:10. Unreacted maleimide dyes were quenched by the addition of 1 mM dithiothreitol (DTT). Subsequently the sample was purified by gel filtration over a S200 10/300GL column from GE Healthcare. Gel filtration Buffer was composed of 25 mM Pipes (pH 6.8), 2 mM MgCl2, 200 mM NaCl, 1 mM EGTA, 1 mM DTT, and 10% sucrose. Finally the sample was flash frozen and stored at −80°C.

### Dynein cloning, purification and labeling

Dynein was expressed and purified as previously described (35). Monomeric constructs for negative stain imaging were further purified by gel filtration on a GE Healthcare Superdex 200 10/300GL in dynein gel filtration Buffer (50 mM K-Ac, 20 mM Tris, pH 8.0, 2 mM Mg(Ac)_2_, 1 mM EGTA, 1 mM TCEP, and 10% glycerol) and flash frozen afterwards. For the negative Sain images we used the VY137 construct with the following genotype: PGal:ZZ:Tev:GFP:HA:D6 MATa; his3-11,15; ura3-1; leu2-3,112; ade2-1; trp1-1; PEP4::HIS5; PRB1D. For the in solution distance measurements we added a c-terminal Halo-tag (28) and inserted a YBBR-tag (29) into the MTBD - VY1067. Before gel filtration, the monomer was labeled on ice overnight with two 16bp long double stranded DNA constructs (D-E and F-G) that were dimerized a priori with Assembly Buffer (20 mM Tris (pH 8.0), 1 mM EDTA, and 2.5 mM MgCl_2_) and heating the mixture to 95°C for 5 min, followed by a cooling of 1°C per minute down to 20°C. The YBBR-tag labeling was carried out as previously described (36). Briefly, we mixed 10 mM MgCl2, 2.5 μM Sfp phosphopantetheinyl transferase, 5 μM DNA–CoA and 50 nM ybbR-tagged dynein (all final concentrations). Afterwards we removed excess DNA strands by gel filtration on a GE Healthcare Superdex 200 10/300GL in dynein gel filtration Buffer (50 mM K-Ac, 20 mM Tris, pH 8.0, 2 mM Mg(Ac)_2_, 1 mM EGTA, 1 mM TCEP, and 10% glycerol) and then flash froze the sample. The oligos were ordered from Biomers GmbH (Ulm, Germany) and Integrated DNA Technologies - IDT - (Skokie, IL) with the following sequences and modifications:

~~~
Strand D:
/CoA/AGGATGAGTGAGAGTG (Biomers)
Strand E: /5BiosG/CACTCTCACTCATCCTT/3Cy3Sp/ (IDT) Strand F:
/HALO/AGGATGAGTGAGAGTG (Biomers)
Strand G: /5BiosG/CACTCTCACTCATCCTT/3ATTO647NN/ (IDT)
~~~

### Preparation of microtubules

Tubulin was purified and polymerized as previously described (37). Unlabeled tubulin, biotinylated tubulin, and fluorescent tubulin were mixed at a ratio of 8:2:1 in BRB80 and 1 mM GTP was added. Then the mixture was incubated in a 37°C water bath for 15 min. Afterwards 20 µM of Taxol (Sigma, T1912) was added and the mixture was incubated for an additional 2 h at 37°C. Before usage, microtubules were spun over a 25% sucrose cushion at ∼160,000 *g* for 10 min in a tabletop centrifuge.

### Preparation of flow-cells with kinesin

Flow-cells with immobilized kinesin were prepared as previously described (17). First, we added 10 µl of 5 mg/ml Biotin-BSA in BRB80 to the flow-cell and incubated for 2 min. Then, we again added 10 µl of 5 mg/ml Biotin-BSA in BRB80 and incubated for 2 min. Afterwards we washed with 20 µl of BRB80 with 2 mg/ml β-casein (Sigma, C6905), 0.4 mg/ml κ-casein (Sigma, C0406). This was followed by addition of 10 µl of 0.5 mg/ml Streptavidin in PBS and a 2 min incubation. We then washed with 20 µl of BRB80 with 2 mg/ml β-casein, and 0.4 mg/ml κ-casein. In a next step 10 µl of polymerized Alexa488 labeled microtubules were added and incubated for 5 min. Next, we washed with 30 µl of BRB80 with 2 mg/ml β-casein, 0.4 mg/ml κ-casein, and 10 µM Taxol. Then, we added 10 µl of K490 in BRB80 with 2 mg/ml β-casein, 0.4 mg/ml κ-casein, 10 µM Taxol, and 1 mM AMPPNP (Sigma, 10102547001) and incubated for 5 min. Afterwards we washed with 30 µl of BRB80 with 1 mg/ml β-casein, 0.2 mg/ml κ-casein, 10 µM Taxol, and 1 mM AMPPNP. Finally we added the PCA/PCD/Trolox oxygen scavenging system (34) in BRB80 with 1 mg/ml *β*-casein, 0.2 mg/ml κ-casein, 10 µM Taxol, and 1 mM AMPPNP.

### Preparation of flow-cells with dynein

The flow cells for the distance measurements between the AAA ring and the microtubule binding domain of dynein were prepared as follows. First, we mixed DNA labeled, monomeric dynein in DAB (50 mM K-Ac, 30 mM HEPES, pH 7.4, 2 mM Mg(Ac)_2_, 1 mM EGTA, 1mM TCEP) with 1 mM Mg-ATP and 1 mM vanadate (Sigma, 450243) and incubated at RT for 15 min. We also prepared a dynein dilution in DAB for the apo state and also incubated it at RT for 15 min. In the meantime we prepared two identical flowcells for the apo and ATP vanadate state on the same microscopy slide. Therefore, we added 10 µl of 5 mg/ml Biotin-BSA in BRB80 twice and incubated for 2 min each time. Afterwards the flowcell was washed with 20 µl of BRB80 with 2 mg/ml β-casein (Sigma, C6905), 0.4 mg/ml κ-casein (Sigma, C0406). We then added 10 µl of 0.5 mg/ml Streptavidin in PBS and incubated for another 2 min. This was followed by a wash with 20 µl DAB with 2 mg/ml β-casein (Sigma, C6905), 0.4 mg/ml κ-casein (Sigma, C0406). Next, we incubated with 10 µl of either dynein solution, apo and ATP vanadate, for 5 min. Afterwards we washed with 30 µl of DAB with 1 mg/ml β-casein, and 0.2 mg/ml κ-casein. For the ATP vanadate state we added 1 mM of Mg-ATP and 1 mM of vanadate. Finally, we added 10 µl of the PCA/PCD/Trolox oxygen scavenging system (34) in DAB with 1 mg/ml β-casein, and 0.2 mg/ml κ-casein. For the ATP vanadate state the Buffer was supplemented with 1 mM Mg-ATP and 1 mM vanadate.

### Microscope setup

Data collection was performed at room temperature (∼23°C) using through-the-objective total internal reflection fluorescence (TIRF) inverted microscopy on a Nikon Eclipse Ti microscope equipped with a 100× (1.45 NA) oil objective (Nikon, Plan Apo λ). We used two stepping motor actuators (Sigma Koki, SGSP-25ACTR-B0) mounted on a KS stage (KS, Model KS-N) and a custom-built cover to reduce noise from air and temperature fluctuations. A reflection based autofocus unit (FocusStat4) was custom adapted to our microscope (Focal Point Inc.). We applied Nikon Type NF2 immersion oil (Nikon, MXA22126) to all slides. Three laser lines at 488 nm (Coherent Sapphire 488 LP, 150 mW), 561 nm (Coherent Sapphire 561 LP, 150 mW), and 640 nm (Coherent CUBE 640-100C, 100 mW) were guided through an AOTF (Neos, 48062-XX-.55), enlarged 6 fold, passed through a quarter wave plate (ThorLabs, AQWP05M-600) and focused using an achromatic doublet f=100 mm on a conjugate back focal plane of the objective outside of the microscope. The TIRF angle was adjusted by moving a mirror and focusing lens simultaneously. A TIRF cube containing excitation filter (Chroma, zet405/491/561/638x), dichroic mirror (zt405/488/561/638rpc), and emission filter (Chroma, zet405/491/561/647m) was mounted in the upper turret of the microscope. The lower turret contained a filter cube (Chroma, TE/Ti2000_Mounted, ET605/70m, T660lpxr, ET700/75m) that directs Cy3 emission towards the back camera and the Cy5 emission towards the left camera. We used two Andor iXon 512×512 EM cameras, DU-897E. The acquisition software was μManager (14) 2.0. All acquisitions were carried out with alternating excitation between the 561 and 640 laser lines (to avoid considerable background fluorescence in the Cy5 channel caused by 561 nm laser excitation). Image pixel size was 159 nm.

### Single-molecule TIRF data collection

For TetraSpeck™ bead acquisitions an exposure time of 100 msec and for all other samples 400 msec was used, if not otherwise specified. After every stage movement we waited 3 sec before collecting data to minimize drift effects. We used the cameras in conventional CCD mode (i.e., no EM gain). All data sets were acquired with a ‘16 bit, conventional, 3 MHz’ setting and a preamp gain of 5x. More details of image acquisition settings and laser powers settings for each individual data set are shown in Table S4, S5, and S6. A step-by-step protocol for data acquisition is given in the SI Protocol.

### Negative stain electron microscopy data collection and processing

Nucleotide-bound samples were prepared with 5 mM ATP + Sodium vanadate in addition to equimolar magnesium acetate. For negative-stain EM, samples were applied to freshly glow discharged carbon coated 400 mesh copper grids and blotted off. Immediately after blotting, a 0.75% uranyl formate solution was applied for staining and blotted off. The stain was applied five times per sample. Samples were allowed to air dry before imaging. Data were acquired at UCSF, on a Tecnai T12 microscope operating at 120 kV, using a 4k×4k CCD camera (UltraScan 4000, Gatan) and a pixel size of 2.1 Å/pixel. Particles were picked and boxed using scripts from SAMUEL and SamViewer (https://liao.hms.harvard.edu/samuel). 2D classification was used to clean our stack and obtain a set of good particles. Only top view (views in which the AAA ring could be clearly identified) were used. Particles were manually scored as having a “full” stalk (MTBD visible), “partial stalk” (stalk is visible but MTBD is not) or “no stalk” (stalk cannot be identified in the micrograph) (Table S1). For an unbiased sorting, we randomly assigned unique identiffiers (10 digid number) to each particle in the apo and ATP-vanadate state, pooled all particles from both nucleotide states, sorted them manually into the three different classes (stalk, partial stalk, no stalk) and then decoded particles based on the unique identiffier to sort the particle back into the apo or ATP-vanadate states.

### Single-molecule localization

All emitters were fitted and localized using μManager’s (14) “Localization Microscopy’ plug-in. For emitter fitting we implemented a Gaussian based maximum-likelihood estimation (15) in μManager (14) and used the following starting conditions. The x- and y-position were determined by centroid calculation, the width was set to 0.9 pixels, background was calculated by summing the intensities of all outermost pixels of an ROI, and intensity was determined by summing up all intensities within the ROI minus the background value. After fitting, intensities and background were converted to photon count by applying the photon conversion factor and correcting for camera offset and read noise. Width and x-, y-coordinates were then converted from pixel to nanometer space (1 pixel = 159 nm). When fitting emitters with μManager’s (14) “Localization Microscopy’ plug-in a noise threshold and box size can be set. Parameters for analysis are shown in Table S4.

We then calculated the variance in fluorophore localization using the MLEwG method (15). Note that we used intensity and background values determined by the aperture method (38) and not values determined by the MLE emitter fitting because the aperture method values agreed better with the experimentally measured variance (Fig. S11). A step-by-step protocol for single-molecule localization is given in SI Protocol.

### Image registration

For image registration two data sets were always acquired: Fiducial markers (TetraSpeck™ beads) to determine the registration map before imaging the sample of interest and a second set of beads to test the stability of the registration (target registration error - TRE) after the sample of interest. Registrations were carried out by first applying a global affine transformation (determined from the bead images) to bring the coordinates in the two channels in close enough proximity for automated pair assignment (Fig. S1). Final registration was accomplished by applying a second affine transform constructed from beads in the immediate vicinity of each pair (i.e., each pair has its own piecewise affine transform). This piecewise affine transformation (9) was also used to calculate the TRE from the second set of bead images by determining the difference in x and y position of each bead after registration (Eq. 1). Since piecewise affine alignment is based on a nearest neighbor search (9), three parameters can influence registration outcome: minimum and maximum number of fiducial points and the maximum sdistance to the control point. Higher maximum distance and higher maximum number of points caused distortions indicating that the registration was not executed properly (Fig. S3). On the other hand, when the maximum distance is too small, an area in the micrograph may not contain the minimum number of fiducials, and thus will not be corrected (white areas in Fig. S3c). Based on the analysis of many different parameter combinations (Fig. S4), we used the following settings for piecewise affine maps: a minimum of 10 and a maximum of 100 fiducial points as well as a maximum distance of 2 μm (except for Fig. S7 where a maximum distance of 3 μm was used, and Fig. S3 where values are provided in the figure caption). A step-by-step protocol for image registration is given in SI Protocol.

### Single-molecule data analysis and distance determination

All data sets were analyzed using μManager’s (14) “Localization Microscopy” plug-in. The fitting method (P2D, Sigma-P2D, Vector-P2D, and Vector) to calculate the distance between two fluorophores is either indicated in the figure and/or figure caption. To avoid erroneous results caused by floating point under- or overflows during the calculation of P2D (3), intermediate results were tested for such conditions and set to minimum or maximum floating point number when appropriate. Furthermore, an approximation (Appendix B of Churchman et al., 2006 (6)) of the P2D function was used

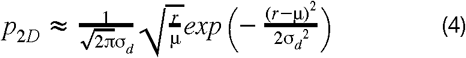

when the estimate of σ_d_ was smaller than half the estimate of the distance.

For P2D and Sigma-P2D the data was fit by means of maximum likelihood estimation (MLE) as described in the results section. For Vector and Vector-P2D we used a more outlier robust fitting method (non-linear least squares (NLLSQ) fitting) since experimental data usually contain some background noise causing incorrect fitting results when using maximum likelihood estimation for Vector-P2D. We could have also removed outliers from the data but it is not always possible to distinguish “real” data points from outliers and small changes in threshold value (cut-off for measured distances) dramatically influence the outcome of the maximum likelihood fit of distance μ. Setting the cut-off for the measured distances too low or too high can dramatically change the value of the estimated distance for MLE fitting. When fitting with NLLSQ, setting the distance cut-off too low might influence the outcome. However, since NLLSQ is less sensitive to outliers, the cut-off can always be set to high values (e.g. 4-5 times of the expected distances) and therewith erroneous fitting results are less likely.

To overcome problems with bin size settings for histograms when fitting with NLLSQ we converted the experimental data into an empirical cumulative distribution function and fit this with numeric integration of the P2D. We show by means of Monte-Carlo simulation that NLLSQ fitting is as good as MLE for data lacking background noise and that NLLSQ fitting outperforms MLE in all conditions where random background noise was added (Fig. S18). The NLLSQ fitting predicts the true distance correctly even for data where 5% of the values are random background noise.

A standard error of the mean (S.E.M.) for distance calculations using Sigma-P2D and P2D (Fig. 3) was determined by means of Fisher Information Matrix whereas bootstrapping was used for Vector-P2D and Vector (Fig. 4 and 5). Parameters for analysis are shown in Table S4 and S6. A step-by-step protocol for data analysis is given in SI Protocol.

### Monte Carlo simulations

*In silico* two-color distance measurements by means of Monte Carlo simulation were carried out with a custom Python script. In brief, the true distance µ, the two localization errors σ_loc_1_ and σ_loc_2_, their underlying distributions (σσ(_loc_1_), σσ(_loc_2_)), sample conformational heterogeneity σ_con_, the number of pairs observed, and the number of frames (observations) per pair can be varied in parallel. The simulation for each parameter combination can be repeated multiple times if desired. For the variance in the fluorophores localization a Gaussian distribution is applied to true positions of channel 1 and 2 and a Gamma distribution is applied as the underlying distribution of the variance in the fluorophores localization for channel 1 and 2. We analyzed model datasets based on different ratios of distance uncertainty to distance (σ_d_ / μ). For each ratio we evaluated 10 datasets with Sigma-P2D and P2D or Vector and Vector-P2D and calculated the average distance to determine the discrepancy between the true distance d and the calculated distance. Based on common localization errors for single-molecule studies (Fig. S10) and distances on the nanometer scale (∼2-30 nm), we expect ratios (σ_d_ / μ) of up to 4 to be of experimental relevance. However, we included even higher ratios to probe the upper limits of Sigma-P2D and Vector-P2D. For more details see SI Software 2.

### Statistics

For each result the inherent uncertainty due to random or systematic errors and their validation are discussed in the relevant sections of the manuscript. Details about the sample size, number of independent calculations, and the determination of error bars in plots are included in the figures and figure captions.

### Code availability

The custom-written Python code for Monte Carlo simulations is available as SI Software published with the online version of the paper under the Berkeley Software Distribution (BSD) license and is hosted on GitHub at: https://github.com/stefanniekamp/localization-microscopy. μManager acquisition and analysis software is available as SI Software partly under the Berkeley Software Distribution (BSD) license, partly under the GNU Lesser General Public License (LGPL) and is hosted on GitHub at https://github.com/nicost/micro-manager. The latest version for MacOS / Windows can be found here: https://valelab.ucsf.edu/∼nico/mm2gamma/

### Data availability

Raw data sets to be used to create and test registration maps, and to measure distances, are provided at https://valelab.ucsf.edu/∼sniekamp/. In addition, we provide a step-by-step SI Protocol that describes how this raw data can be analyzed with μManager’s (14) ‘Localization Microscopy’ plug-in. The raw electron microscopy micrographs can be found at https://valelab.ucsf.edu/∼sniekamp/. All other data files are available upon request.

## ACKNOWLEDGMENTS

We thank A. Jain, D. Larsen, and T. Skokan for critical discussions of the manuscript. We are grateful to E. Jonsson for the kinesin plasmids and discussions about the kinesin distance measurements. The authors acknowledge funding from the National Institutes of Health (R01EB007187, R.D.V. and 1F32GM113366-01, J.S., and R00GM112982, G.B.), the Damon Runyon Cancer Research Foundation DFS-20-16 (G.B.) and the Howard Hughes Medical Institute.

## AUTHOR CONTRIBUTIONS

S.N., J.S., G.B., R.D.V., and N.S. designed the research; N.S. and S.N. developed the software; S.N. and J.S. performed Monte Carlo simulations; S.N., W.H., and G.B. prepared samples; S.N. collected TIRF microscopy data; G.B. collected electron microscopy data; S.N., J.S., G.B., and N.S. analyzed the data; S.N., R.D.V., and N.S. wrote the article; All authors read and commented on the paper.

